# Construction of an Immunoinformatics-Based Multi-Epitope Vaccine Candidate targeting Kyasanur Forest Disease Virus

**DOI:** 10.1101/2024.03.14.584963

**Authors:** Sunitha M. Kasibhatla, Lekshmi S. Rajan, Anita M. Shete, Vinod Jani, Savita Patil, Yash Joshi, Rima R. Sahay, Deepak Y. Patil, Sreelekshmy Mohandas, Triparna Majumdar, Uddhavesh Sonavane, Rajendra Joshi, Pragya D. Yadav

**Author notes:** Correspondence: Dr. Pragya D. Yadav, Scientist F & Head, Maximum Containment Facility, ICMR-National Institute of Virology, Pune, Maharashtra, India, Dr. Rajendra Joshi, Senior Director and HOD, HPC-Medical and Bioinformatics Applications Group, Centre for Development of Advanced Computing, Pune University Campus, Ganesh Khind, Pune - 411 007, Maharashtra, India. Equal contribution as first author (Sunitha M. Kasibhatla, Lekshmi S. Rajan, Anita M. Shete).

## Abstract

Kyasanur Forest Disease (KFD) is one of the neglected tick-borne viral zoonoses. KFD virus was initially considered endemic to the Western Ghats region of Karnataka. Still, over the years, there have been reports of its spread to newer areas within and outside Karnataka. The absence of an effective treatment for KFD expedites the need for further research and development of novel vaccines. The present study was designed to develop a multi-epitope vaccine candidate against KFDV using immunoinformatic tools. After analyzing 74 complete KFDV genome sequences for genetic recombination and phylogeny, different prioritized B and T cell epitopes were combined using various linkers to construct the vaccine candidate. Docking analysis of the designed vaccine construct revealed a stable interaction with the TLR2-TLR6 receptor complex. After confirming the stability of the vaccine receptor complex, codon optimization was done to ensure the efficient translation of the designed multi-epitope vaccine in the prokaryotic host system, and the subsequent *in-silico* cloning into the pET30b(+) expression vector was carried out. Immunoinformatics analysis of the multi-epitope vaccine in the current study is satisfactory as it can significantly accelerate the initial stages of vaccine development by narrowing down potential vaccine candidates and providing insights into their design. Experimental validation of the potential multi-epitope vaccine candidate remains crucial to confirm effectiveness and safety in real-world conditions.

## 1. Introduction

Kyasanur Forest Disease is a highly neglected emerging tick-borne viral zoonosis caused by Kyasanur Forest Disease Virus (KFDV) that belongs to the family *Flaviviridae*. The disease was first identified in 1957 in the Kyasanur forest region of Karnataka, India, and has since posed a significant public health threat, with sporadic outbreaks occurring in newer districts of Karnataka and the states of Maharashtra, Goa, Kerala, and Tamil Nadu^1^. The mortality rate for KFDV infection is reported to be about 2–10%^2^. The virus is maintained in mammals and birds and is transmitted through ticks to the vertebrate hosts, with the main vector being the anthropophagic *Haemaphysalis spinigera*^3–5^.

KFDV is an enveloped spherical virus with single-stranded RNA of 11kb size in the icosahedral nucleocapsid. The virion has a 40-65 nm diameter and codes for a single polyprotein^6,7^. The envelope protein of KFDV plays an important role in the entry of the pathogen into the host cells by receptor binding and membrane fusion^8–9^. The dimeric and trimeric forms of E protein play a role in the post-fusion stability of the virion. It evokes the host immune response, produces neutralizing antibodies, and is considered a suitable candidate for developing new vaccines and diagnostics^10^.

Symptomatic treatment and supportive therapy are followed to treat the affected individuals, as there is no clinically approved drug against KFDV. The formalin-inactivated KFD vaccine was one of the early approaches to vaccinating against the disease and has been used primarily in certain regions of India where KFD is endemic. However, the primary concern with the formalin-inactivated vaccine is its hesitancy by people due to side effects at the injection site^11,12^. While it may offer some protection against KFDV, it may not provide complete or long-lasting immunity. There may be breakthrough infections among individuals who have received the vaccine. Earlier studies demonstrated the low effectiveness of the vaccine even after repeated booster doses^13,14^. Besides this, the estimated mean rate of nucleotide substitution based on the complete genome of KFDV is 4.2 × 10^−4^ subs/site/year^15^. This means that, on average, one nucleotide in the KFDV genome is expected to change every 23 years. The earliest known KFDV strain was isolated in 1957, so the virus has evolved for at least 66 years. Studies have shown that KFDV can accumulate mutations in its envelope protein. Some of these mutations have been linked to changes in the antigenicity of the envelope protein, which means that the virus may become less susceptible to the immune system^15–17^. The low efficacy of currently available vaccines and the increasing spread of KFDV to naive regions in India necessitates the need for the development of a new KFD vaccine^1,13,14^.

Newer vaccine development techniques, such as recombinant DNA technology or viral vector vaccines, may offer advantages in terms of efficacy and safety. Computational vaccinology and *in-silico* prediction of the host’s immune response to pathogens/antigens can expedite the development of novel vaccines. Epitope-based vaccines, which target specific antigenic regions of the pathogen, offer a promising approach to vaccine development termed ‘reverse vaccinology’ due to their potential for inducing targeted immune responses while minimizing side effects^18^. Computational epitope prediction and immuno-informatics can hasten the design and development of multi-epitope vaccines. The molecular docking and molecular simulations may help in understanding the binding potential of vaccine construction with host immune cell receptors^19^. Immunoinformatics has been gainfully employed for designing multi-epitope vaccine candidates against KFDV (with a relatively smaller set of genome sequences) and to understand the molecular interactions between the host receptors and virus protein recently^20–22^. The present work is a comprehensive phylogenomic and immunoinformatics analysis of KFDV based on 74 genome sequences to rationally design a multi-epitope vaccine candidate based on a major antigenic protein, envelope protein.

## 2. Materials and Methods

### 2.1 Study design

The complete genome sequences of the KFDV isolates available with the ICMR-National Institute of Virology, Pune, India were analyzed using different virus bioinformatics tools to determine the lineages and the level of genetic recombination among the isolates. The study aims to design a vaccine candidate that can induce a strong and specific immune response by targeting the envelope protein that plays a key role in the virus-host interaction. However, it is also important to consider the diversity of KFDV strains and the potential for antigenic variation when designing vaccines based on the envelope protein to ensure broad protection. Considering this, various immunoinformatics tools were used in the current study to create an *in-silico*-designed multi-epitope vaccine directed toward the envelope protein of KFDV. Prioritized B- and T-cell epitopes from the envelope protein of KFDV were used for the vaccine design. The physicochemical properties of the vaccine construct were evaluated for the prediction of the three-dimensional (3D) structure. To analyze the binding affinity and stability, molecular docking analysis and molecular dynamics simulation were performed respectively. **Figure 1** shows the different steps for designing the multi-epitope vaccine construct for KFDV in our study.

**Figure 1:**
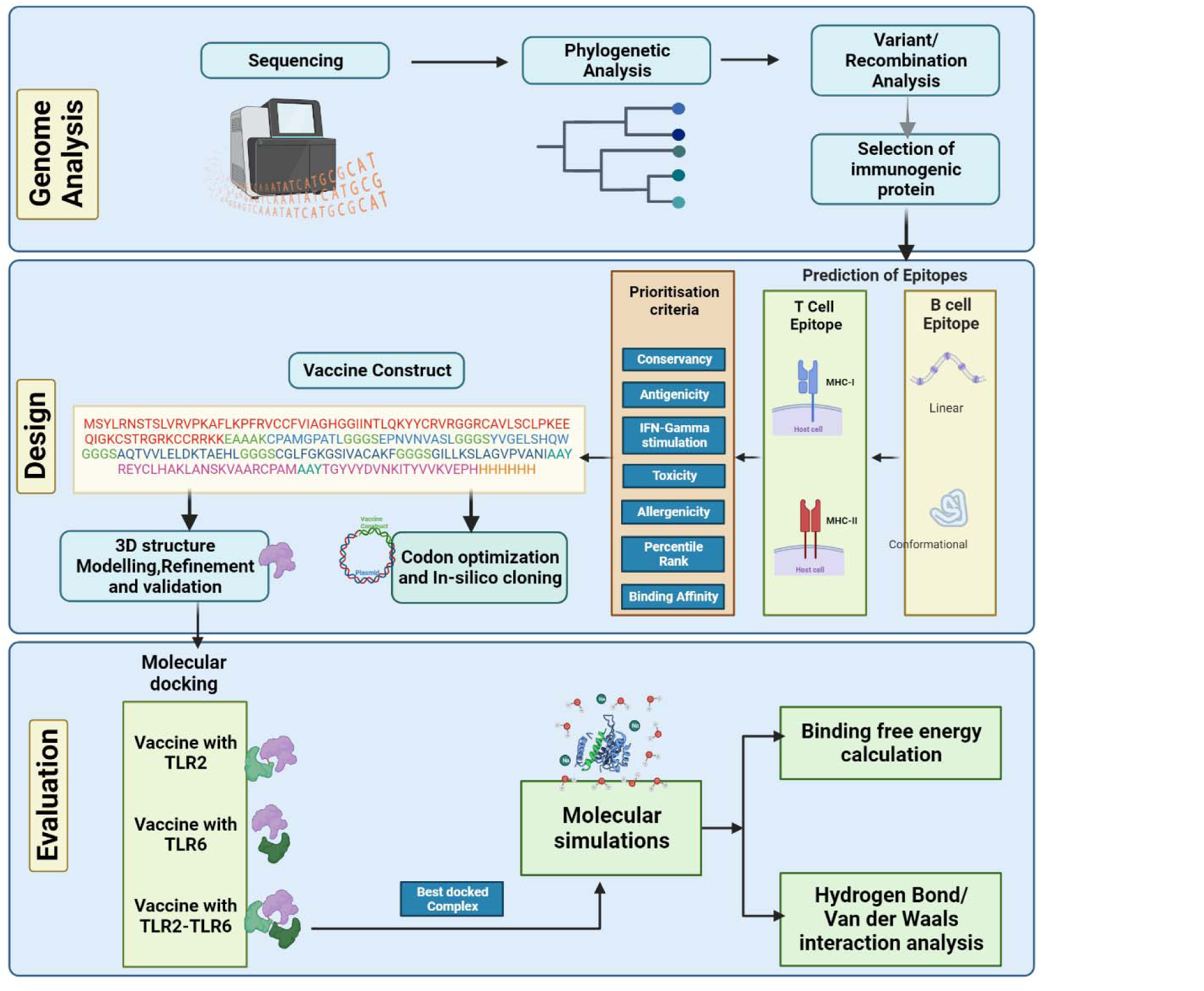
Schematic representation of steps involved in the in silico design of Kyasanur Forest Disease vaccine

### 2.2 Next-generation Sequencing and Phylogenetic Analysis

A total of 74 complete genome sequences of KFDV from India were used in this study. Out of the 74 sequences, (n=50) were previously submitted to NCBI by ICMR-NIV Pune, and (n=24) collected during (2019-2022) were recently sequenced by the Next generation sequencing approach. For Next Generation Sequencing, RNA was extracted from isolates and clinical specimens with a commercially available RNA extraction kit (MagMAX™ Viral/Pathogen, Themofisher Scientific, MA, USA). The libraries were prepared, pooled, and loaded onto the Illumina platform as per the protocol described earlier^23^. CLC Genomics software 22.0.2 (Qiagen, USA) was used for the sequence analysis. Reference mapping was carried out with the KFDV reference genome (Ref Seq: NC_039218) to retrieve the complete genome. Individual sequences were aligned with the reference sequence using MAFFT^24^. Multiple genome alignment and individual genes/proteins were also carried out using MAFFT. MEGAX was used for the visualization of genome/gene/protein alignments^25^. Recombination events were detected using RDP4 with *p-*value ≤0.005. A sequence is said to be recombinant if predicted by at least three different methods available in RDP4 package^26^. The best nucleotide substitution model was estimated by ModelFinder^27^. For the construction of the phylogenetic tree, the maximum likelihood method was implemented in IQTREE^28^ and visualized using iTOL^29^. Molecular clock behavior was tested using TempEst^30^. BEAST v.10.1.4 was used for the estimation of nucleotide substitution rate and lineage divergence^31^. One billion steps of Markov Chain Monte Carlo (MCMC) were carried out in triplicate. The maximum clade credibility tree was visualized using FigTree.

### 2.3 Immunoinformatic analysis

#### 2.3.1 T cell epitopes prediction

NetMHCpan EL 4.1 and NetMHCIIpan 3.2 from the IEDB server were used respectively for predicting MHC I and MHC II epitopes^32^. MHC I and MHC II epitope predictions were carried out with HLA alleles as curated by Weiskopf et al. 2012^33^ and Greenbaum J. et al. 2011^34^, respectively. Further filtration of the predicted epitopes was based on MHC binding affinity, percentile rank and by removing largely overlapping epitopes that share the same core region.

#### 2.3.2 B-cell epitope prediction

Predication of Conformational and Linear B-cell epitopes was done using Bepipred and ElliPro online tools ^35–36^. For the conservancy of the epitopes in the global isolates, the Epitope Conservancy Analysis tool (IEDB server) was used ^37^. IFN-gamma stimulation of the predicted epitopes was predicted using IFNepitope^38^. Further prioritization of the predicted epitopes was carried out by testing for antigenicity using Vaxijen 2.0 [10.1186/1471-2105-8-4]. Allergenicity and toxicity testing were done using AllerTopv2.0 and ToxinPred^39,40^, respectively.

#### 2.3.3 Construction of multi-epitope vaccine

The selected T-cell and B-cell epitopes were conjugated with different linkers. The linker regions used were EAAAK, GGGS, and AAY. The N-terminal of the vaccine construct was linked with an adjuvant β-defensin by the EAAAK linker. MHC I and MHC II epitopes were linked by GGGS and the B cell epitopes by AAY. A histidine tag was included at the C terminal for further purification. The use of adjuvant β-defensin would help to enhance the vaccine immunogenicity^41^.

#### 2.3.4 Physiochemical properties

Various physiochemical properties of the vaccine construct were calculated using the online web server ProtParam (https://web.expasy.org/protparam)

#### 2.3.5. Prediction of 3D structure, refinement, and validation

The three-dimensional tertiary structure of the vaccine construct was determined by the I-TASSER webserver^42^. I-TASSER stands for Iterative Threading Assembly Refinement. The I-TASSER develops multiple models, and based on the C-score, the best model is chosen. The chosen vaccine construct structure was further refined using molecular dynamics simulation using the AMBER20 simulation package^43^ and amberff14SB force field. The system preparation was done where the structure was solvated in a water box where the TIP3P model represented water. The system was neutralized by adding charged ions. The structure was minimized in two stages, the first steepest descent minimization followed by conjugate gradient minimization. Minimization was followed by temperature ramping up to 300K. This was followed by an equilibration at NPT condition for one ns, and finally, a production run of 50 ns was carried out. The final model obtained from the simulation was validated by Ramachandran plot analysis using the Procheck webserver.

### 2.4 Molecular docking with immune cell receptors

The vaccine construct must interact with the immune cell receptors to evoke an immune response. Hence, the binding affinity of the TLR2, TLR6, and TLR2-TLR6 receptor complex with the vaccine construct was studied using the molecular docking technique. HADDOCK2.4 webserver^44^ was used for docking studies with TLR2 receptor (PDB ID: 2z7x), TLR6, and TLR2-TLR6 receptor complex (PDB ID: 3a79) ^44,45^. Prior to docking, the active and passive residues were predicted using WHISCY [https://wenmr.science.uu.nl/whiscy/]. HADDOCK provides multiple docked cluster models. Based on the lowest HADDOCK score, the best-docked cluster was chosen and subjected to molecular dynamics simulations.

### 2.5 Molecular simulations of TLR2-TLR6-vaccine construct

Molecular dynamics simulations were performed for the docked TLR2-TLR6-vaccine construct complex using the AMBER20 simulation package. A similar protocol to that of the vaccine construct was followed for the simulations, except the production run of 240 ns was performed. Interactions analysis and other statistical analysis were performed on the last 200 ns of simulation data. Root Mean Square Deviation (RMSD), Root Mean Square Fluctuation (RMSF), and hydrogen bond analysis were performed using AmberTools21^43^. Interaction calculations were performed using the GetContact tool [https://getcontacts.github.io/] and the LIGPLOT tool^46^. Free energy of binding between the vaccine construct and TLR2-TLR6 receptor, vaccine construct, and TLR2 receptor, and vaccine construct and TLR6 receptor were computed using the MM-GBSA module of the AMBER20 package.

### 2.6 Codon optimization and In silico cloning

Reverse translation and codon optimization was performed using VectorBuilder (https://en.vectorbuilder.com/tool/codon-optimization.html) to have a high protein expression level in *E.coli* (strain K12). The codon-optimized sequence was cloned into pET30b(+) using XhoI and NdeI (https://www.snapgene.com/).

## 3. Results

### 3.1 Analysis of KFDV genomes

There was no evidence of recombination in the KFDV isolates. A linear relationship between root-to-tip distance and time of isolation was observed indicating molecular clock behavior in the dataset **(Figure 2)**

**Figure 2:**
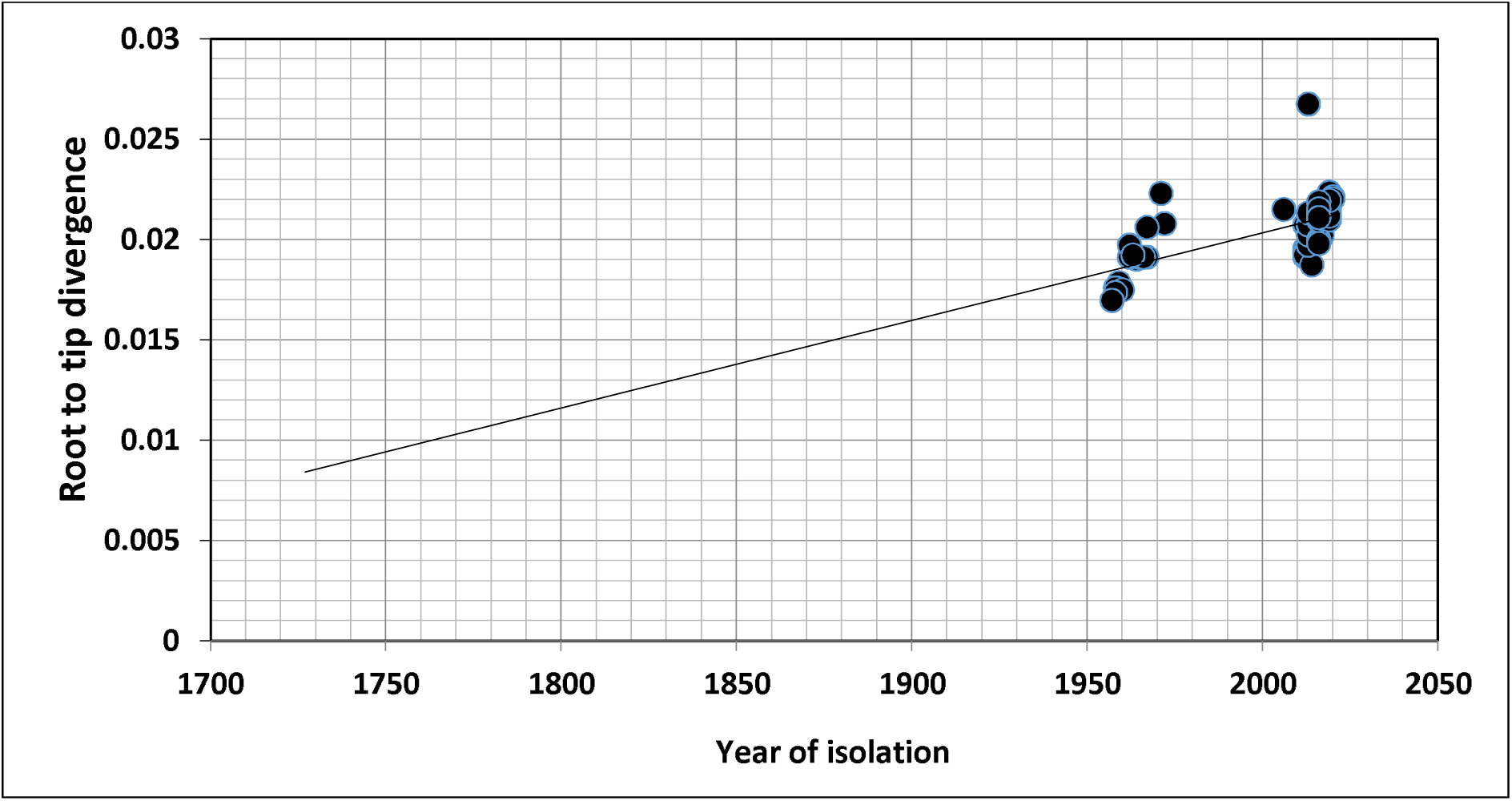
Root-to-tip divergence of KFDV isolates based on complete genome

### 3.2 Phylogenetic analysis

Genome-wide nucleotide substitution rate was found to be 4.31×10^-4^ (95% HPD: 3.45×10^-4^, 5.22×10^-4^) substitutions per site per year. Phylogenetic analysis revealed spatio-temporal clustering with few deviations **(Figure 3).** Two distinct lineages of KFDV were found, of which, lineage 1 includes isolates sampled from Karnataka during 1957-1972. Lineage 2 was found to demarcate into sub-lineages viz., 2.1 and 2.2. Sub-lineage 2.1 includes isolates from Karnataka and Goa during 2006-2022. Sub-lineage 2.2 further differentiates into 2.2.1 and 2.2.2. It was observed that ten KFDV isolates from Karnataka sampled during 2012-2022 constitute lineage 2.2.1. KFDV strains isolated during 2013-2020 from Maharashtra along with six from Karnataka, two from Goa, and one from Tamil Nadu constitute sub-lineage 2.2.2. Nine representative isolates were identified based on the lineages observed in the phylogenetic tree **(Supplementary table 1).**

**Figure 3:**
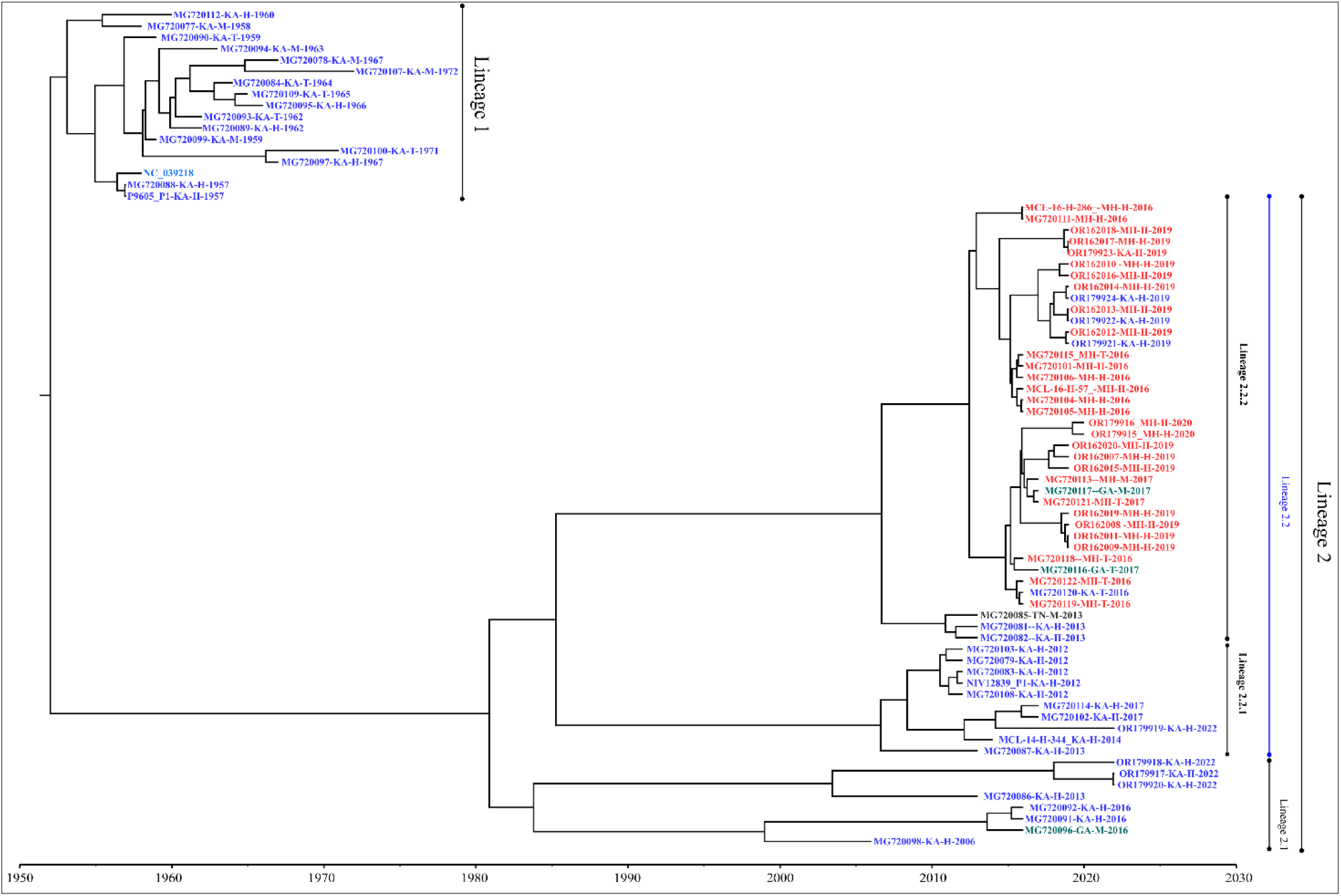
Maximum clade credibility tree of KFDV derived using complete genome

### 3.3 T-cell epitope prediction

MHC class I epitope prediction for envelope protein was carried out for reference HLA alleles as curated by Weiskopf et al. 2012^33^. The predicted epitopes were further filtered based on percentile rank ≤ 1 and predicted binding affinity of 500 nM which resulted in a total of 23 MHC class I epitopes **(Table 1).** Similarly, MHC class II epitopes were predicted for the HLA reference set curated by Greenbaum J. et al. 2011^34^. The class II epitopes were prioritized based on removing largely overlapping epitopes that shar the same core region, MHC binding affinity IC50 ≤ 1000nM, and percentile rank ≤ 10.0 which resulted in a total of 15 MHC class II epitopes **(Table 2).**

**Table 1:**
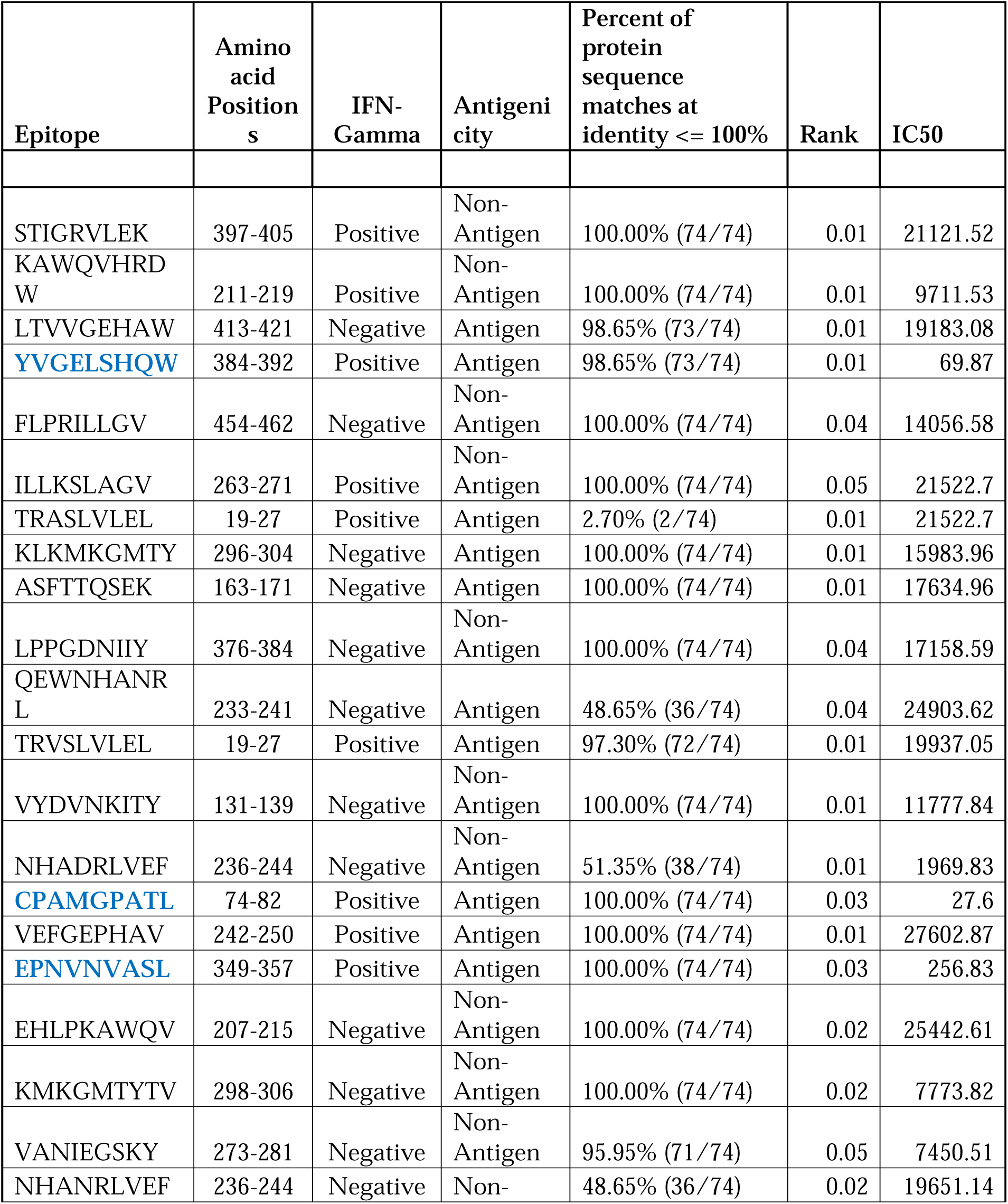
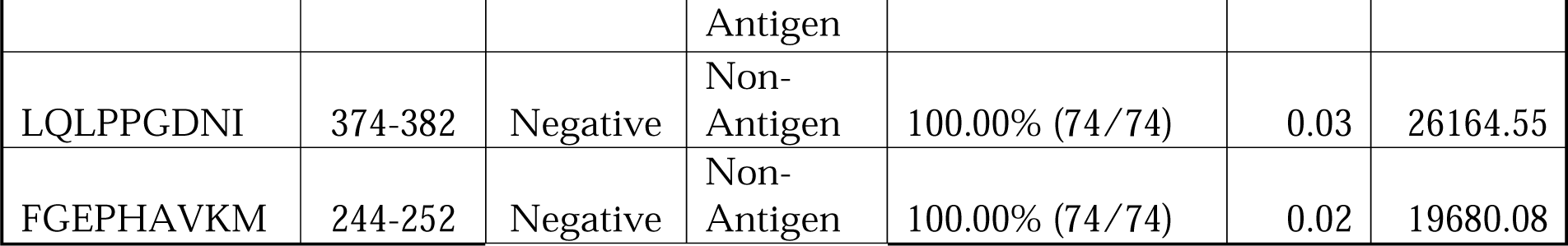
List of short-listed MHC-Class I epitopes belonging to the envelope protein of KFDV. Epitopes used in the vaccine construct are highlighted in bold blue font.

**Table 2:**
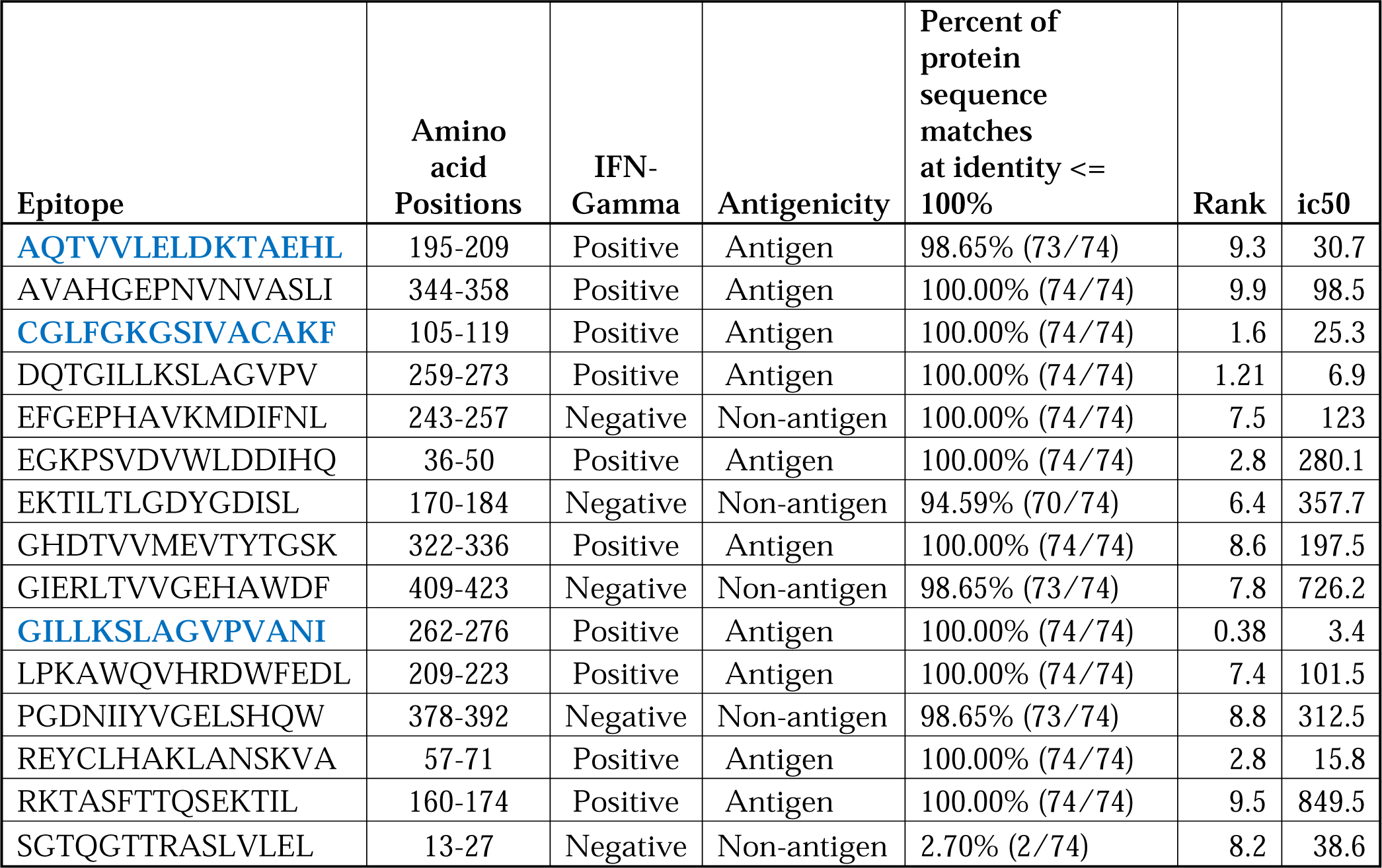
List of short-listed MHC-Class II epitopes belonging to the envelope protein of KFDV. Epitopes used in the vaccine construct are highlighted in bold blue font.

### 3.4 B-cell epitope prediction

Linear B-cell epitopes were predicted using the envelope protein sequences of the nine representative KFDV isolates which were further prioritized based on their conservancy across all global KFDV isolates, the ability to induce IFN-Gamma stimulation, and antigenicity **(Table 3).** The predicted 3D structure of envelope protein was used for conformational epitope prediction and three epitopes were prioritized based on similar criteria as described above for linear B-cell epitopes **(Table 4).**

**Table 3:**
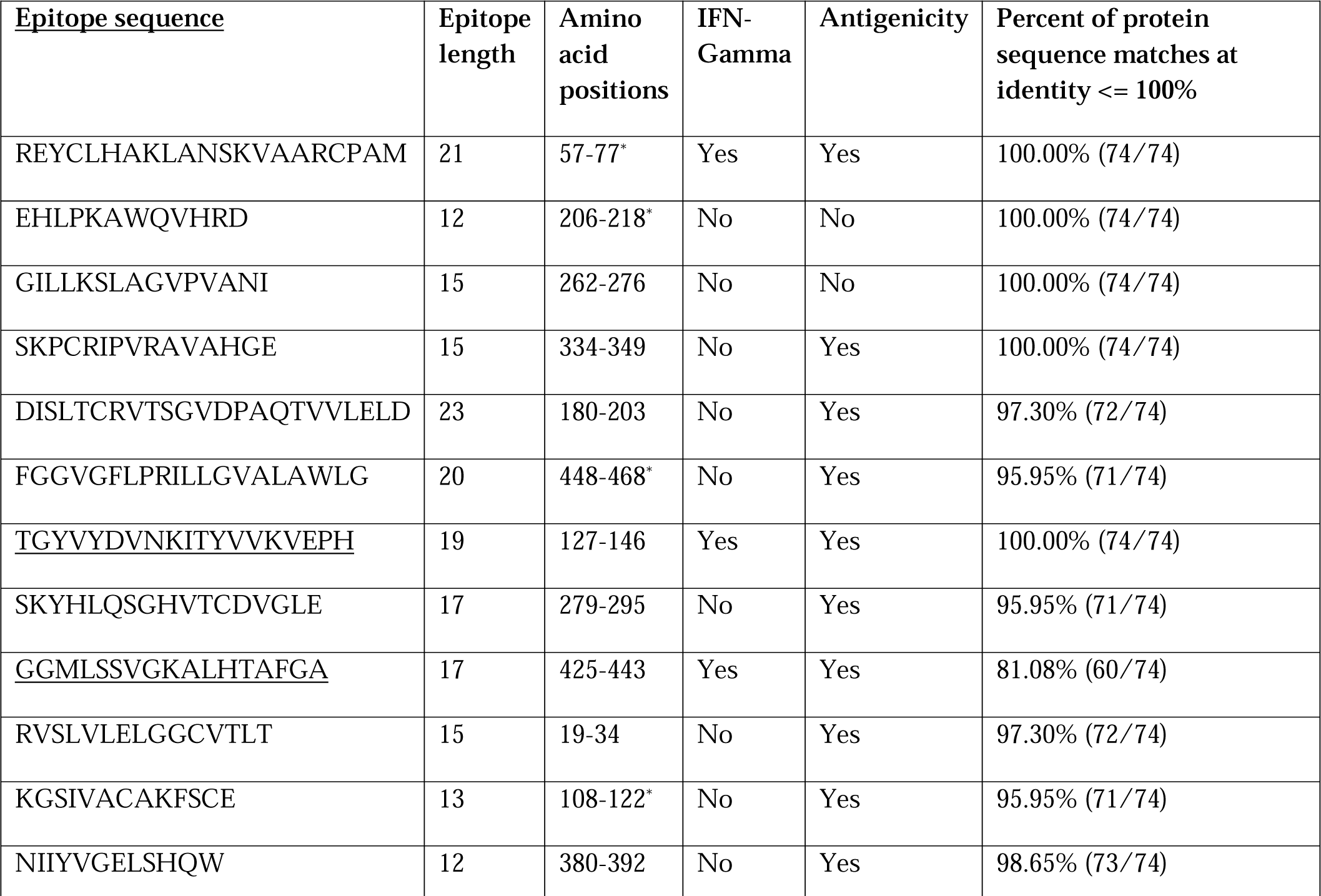
List of predicted linear B-cell epitopes belonging to the envelope protein of KFDV. Prioritized epitopes are underlined. Asterisk indicates experimentally validated epitopes in other *Flavivirus* members47.

**Table 4:**
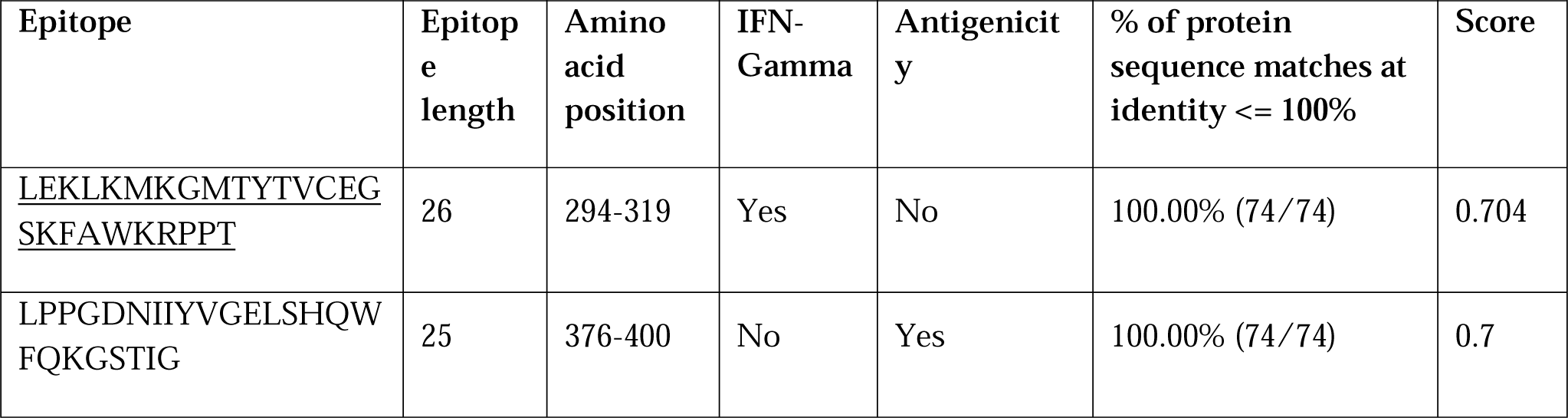
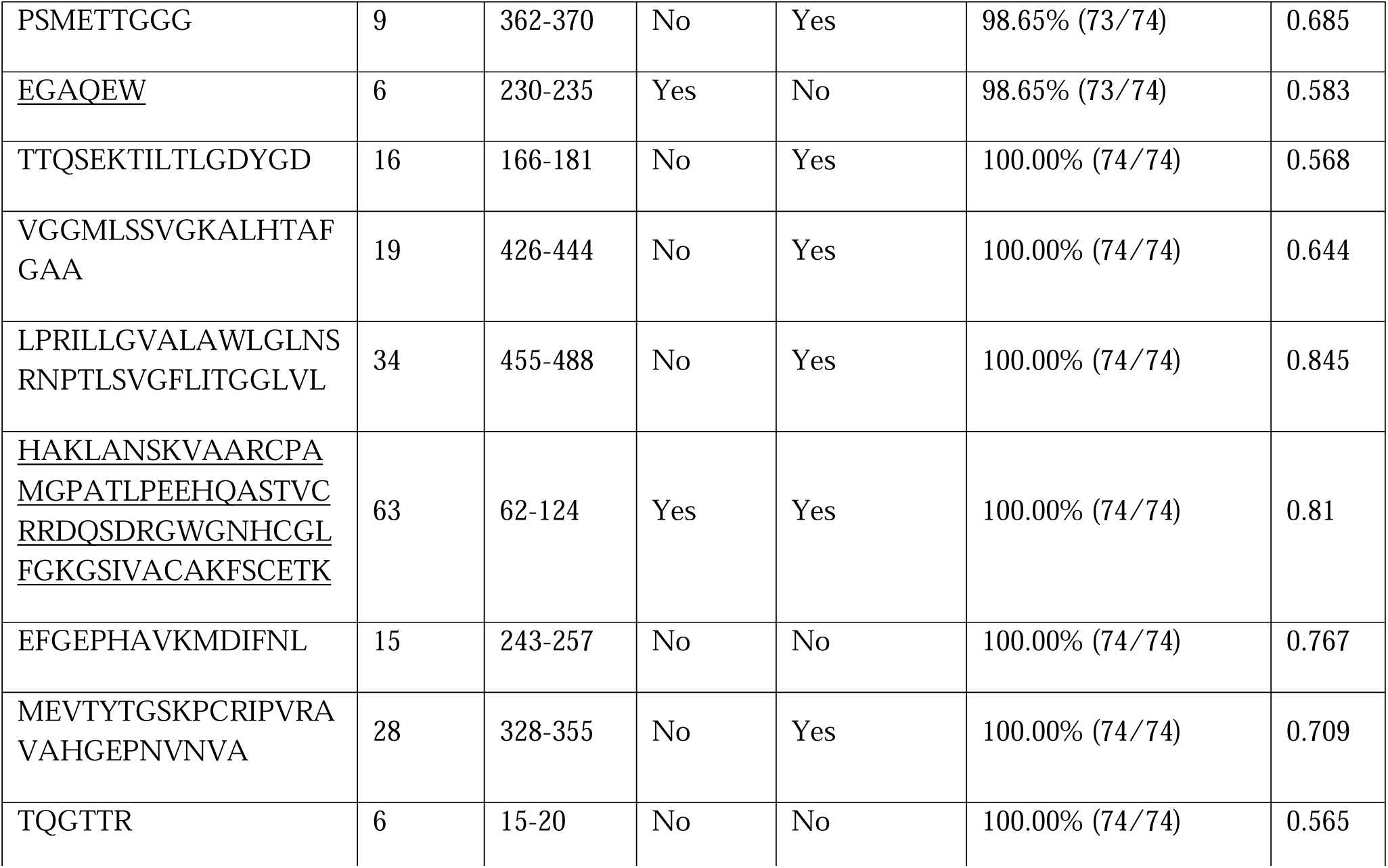
List of predicted conformational B-cell epitopes belonging to the envelope protein of KFDV. Prioritized epitopes are underlined.

**Table 5:** Table representing the statistical parameters for interaction between vaccine construct with TLR2, TLR6, and TLR2-TLR6 construct.

|  | **TLR2** | **TLR6** | **TLR2-TLR6** |
| --- | --- | --- | --- |
| HADDOCK score | -20.2 +/- 3.8 | -82.8 +/- 37.2 | -13.0 +/- 9.3 |
| Cluster size | 19 | 9 | 9 |
| RMSD from the overall lowest-energy structure | 0.8 +/- 0.1 | 1.4 +/- 1.0 | 0.6 +/- 0.2 |
| Van der Waal's energy | -124.0 +/- 19.1 | -146.2 +/- 11.1 | -117.8 +/- 14.8 |
| Electrostatic energy | -302.0 +/- 49.0 | -501.1 +/- 63.6 | -526.7 +/- 32.8 |
| Desolvation energy | -36.8 +/- 6.2 | -10.9 +/- 3.8 | -9.2 +/- 6.2 |
| Restraints violation energy | 2009.5 +/- 219.22 | 1745.3 +/- 162.66 | 6996.5 +/- 225.43 |
| Buried Surface Area | 3996.0 +/- 365.4 | 4874.8 +/- 323.6 | 4564.2 +/- 299.3 |
| Z-Score | -2.3 | -2.1 | -2 |

### 3.5 Construction of multi-epitope vaccine candidate

The prioritized epitopes were combined with the help of different linkers like EAAAK, GGGS, and AAY to construct the linear vaccine. The β-defensin adjuvant was connected to the N-terminal of the vaccine construct using EAAAK linker to boost the immune response. A histidine tag was added to the C-terminal of the vaccine construct to facilitate the vaccine purification. The length of the final vaccine construct is 226 amino acids with a molecular weight of 23.09 kDa **(Figure 4).** The vaccine construct has a theoretical pI value of 9.54, suggesting its alkaline nature.

**Figure 4:**
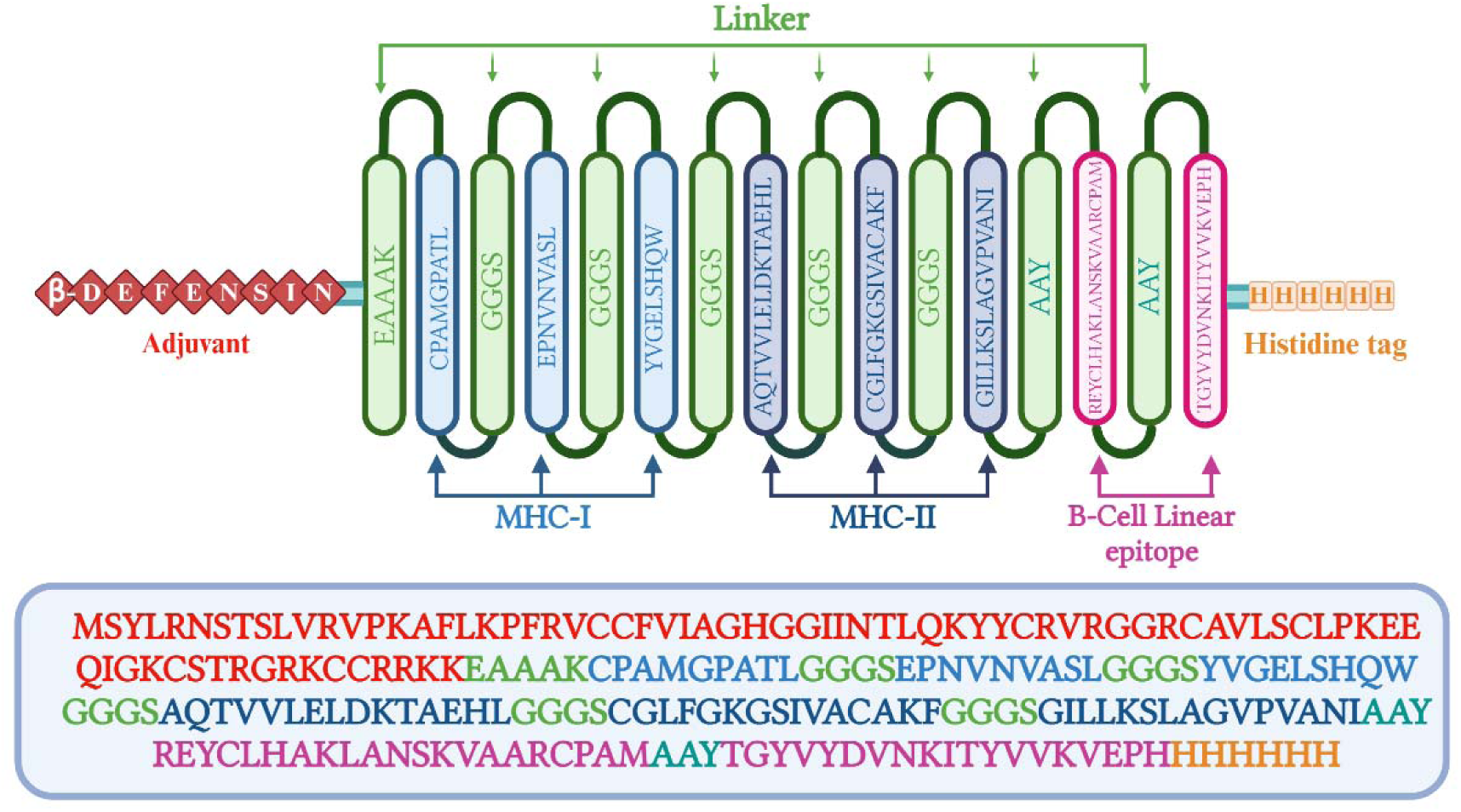
Schematic representation of multi-epitope vaccine with adjuvant at the N-terminal and His-tag at the C-terminal.

The computed instability index (II) was found to be 39.08 which classifies the protein as stable. The average hydropathicity was around -0.029, indicating the hydrophilic nature of the vaccine construct.

### 3.6 Prediction of 3D structure, refinement, and validation

For the prediction of tertiary structure, the I-TASSER server was used. The server predicted five models, and based on the C-score, the top-scoring model was selected for further refinement. The structure was refined with the help of molecular dynamic simulations. The structural quality of the predicted and refined vaccine 3D structure was validated by the Ramachandran plot **(Supplementary** Figure 1). ∼ 98.4% of amino acids belong to the most favored, additional allowed, and generously allowed regions. Three residues, i.e., 1.6 %, belonged to the disallowed region. A good quality predicted structure was predicted with a Z score of -3.54 using ProSA analysis.

### 3.7 Docking of vaccine construct with host receptors (TLR2, TLR6, and TLR2-TLR6 complex)

The docking of host receptors (TLR2-TLR6 complex) with vaccine construct **(Figure 5)** resulted in 73 structures that group into 10 cluster(s). These represent 36.5% of the generated water-refined models. The top cluster was considered the most reliable based on the HADDOCK score. Further refinement of a representative model of the top cluster was carried out using the HADDOCK refinement server. The nine structures in the cluster grouped, resulting in 100% of the water-refined models. T**able 5** shows the statistics for the refined model with TLR2, TLR6, and TLR2-TLR6 complex. The proximity between the vaccine construct and TLR receptors is indicated by a buried surface area of 4564.2 +/- 299.3 Å^2^. The RMSD of the docked complex is around 0.6 +/- 0.2, further reiterating the model’s good quality.

**Figure 5:**
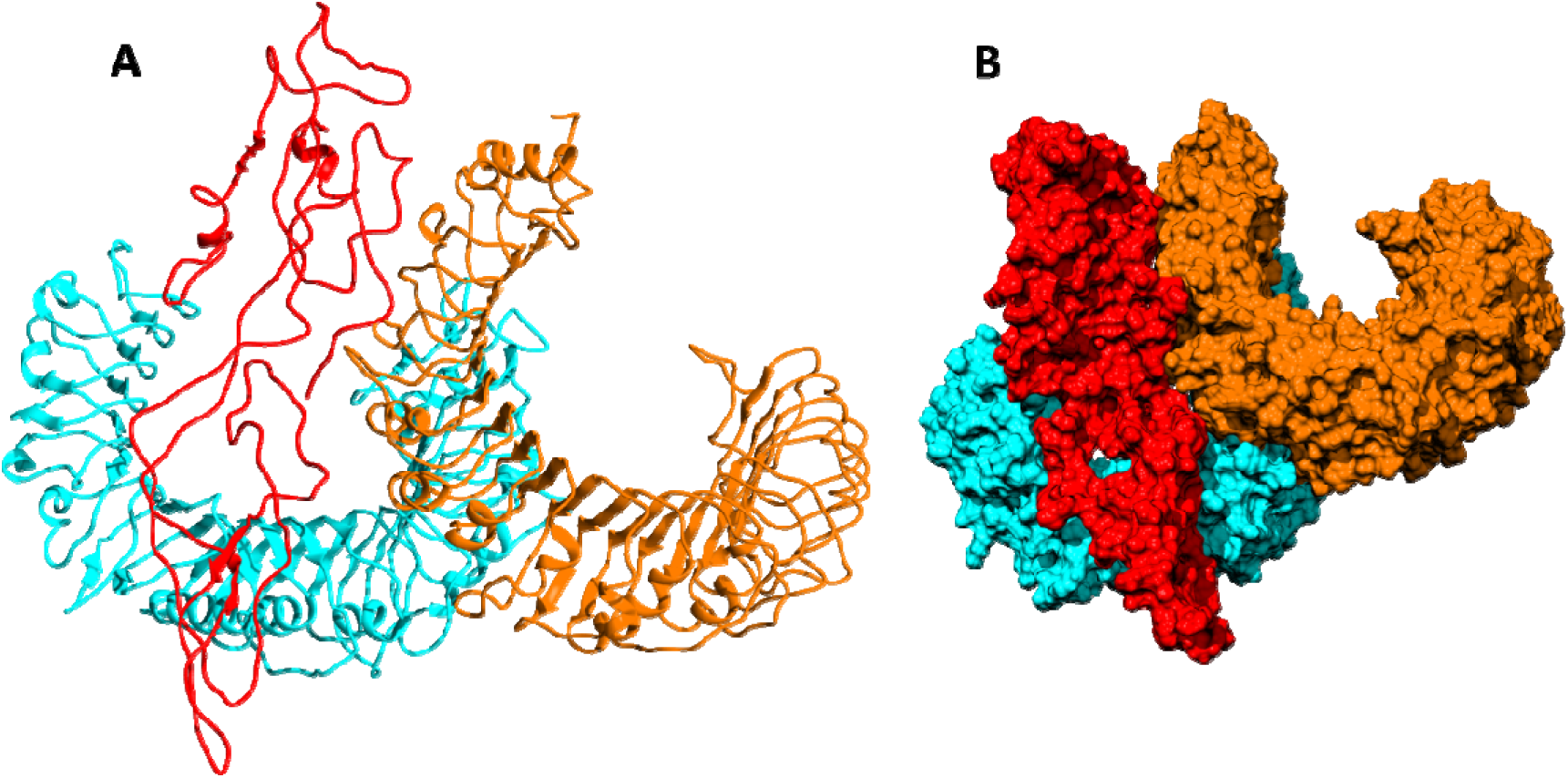
Docked structure of vaccine construct with human TLR2-TLR6. **A**) Two-D structure of the docked model, **B**) Three-D structure of the docked model. The red color represents the vaccine construct, the blue color TLR6, and the orange color TLR2

### 3.8 Molecular simulations of TLR2-TLR6-vaccine construct

Molecular simulations were carried out for the top-scored dock model. Simulation for a total length of ∼240 ns was carried out. The last 200 ns of the molecular simulation was considered for the various structural analyses. RMSD was computed, which indicates how much the structure has deviated from the reference structure. The reference structure considered for the RMSD calculation is the start structure of the simulation. RMSD for the entire complex and individual proteins, i.e., TLR2, TLR6 receptors, and vaccine construct, were computed (**Figure 6).** The RMSD value for the entire protein lies in the range of 4 to 5 Å. For the TLR2 and TLR6 receptors, it was in the range of 2 to 3 Å, and for the vaccine construct, it was in the range of 6 to 7 Å. This clearly shows that the RMSD values tend to stabilize over time, indicating equilibration of the system. The higher RMSD value in vaccine construct may be attributed to the loop regions. Further, to inspect which regions of the vaccine construct show higher fluctuations, RMSF was calculated for the vaccine construct **(Figure 7).** From **Figure 7**, it can be observed that the region with a residue range of 80 -100 and a residue range of 130 - 150 show higher fluctuations.

**Figure 6:**
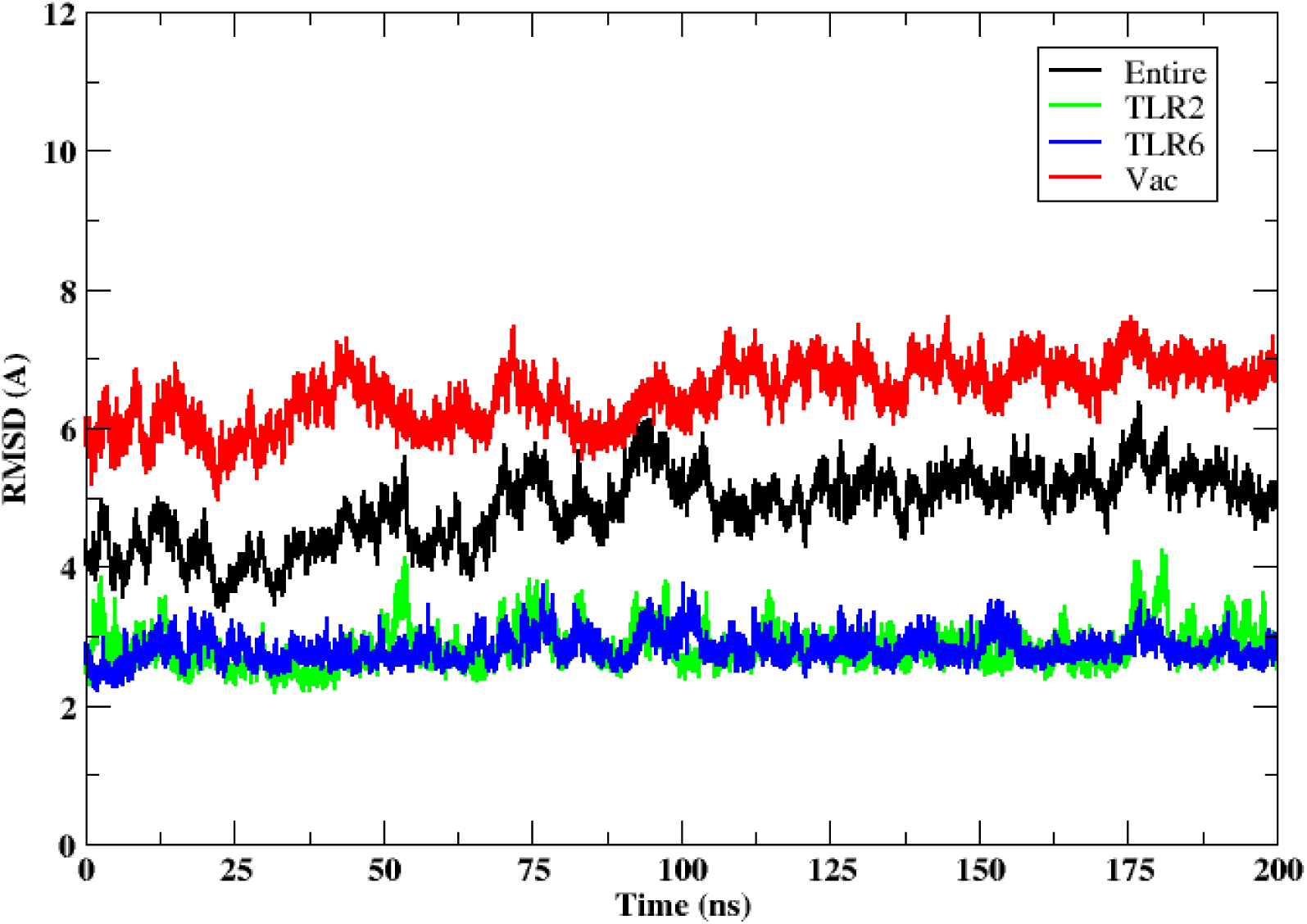
RMSD of the docked 3D structure of vaccine construct with TLR2-TLR6. RMSD of the entire complex (black), vaccine construct (red), TLR2 receptor (green) and TLR6 receptor (blue).

**Figure 7:**
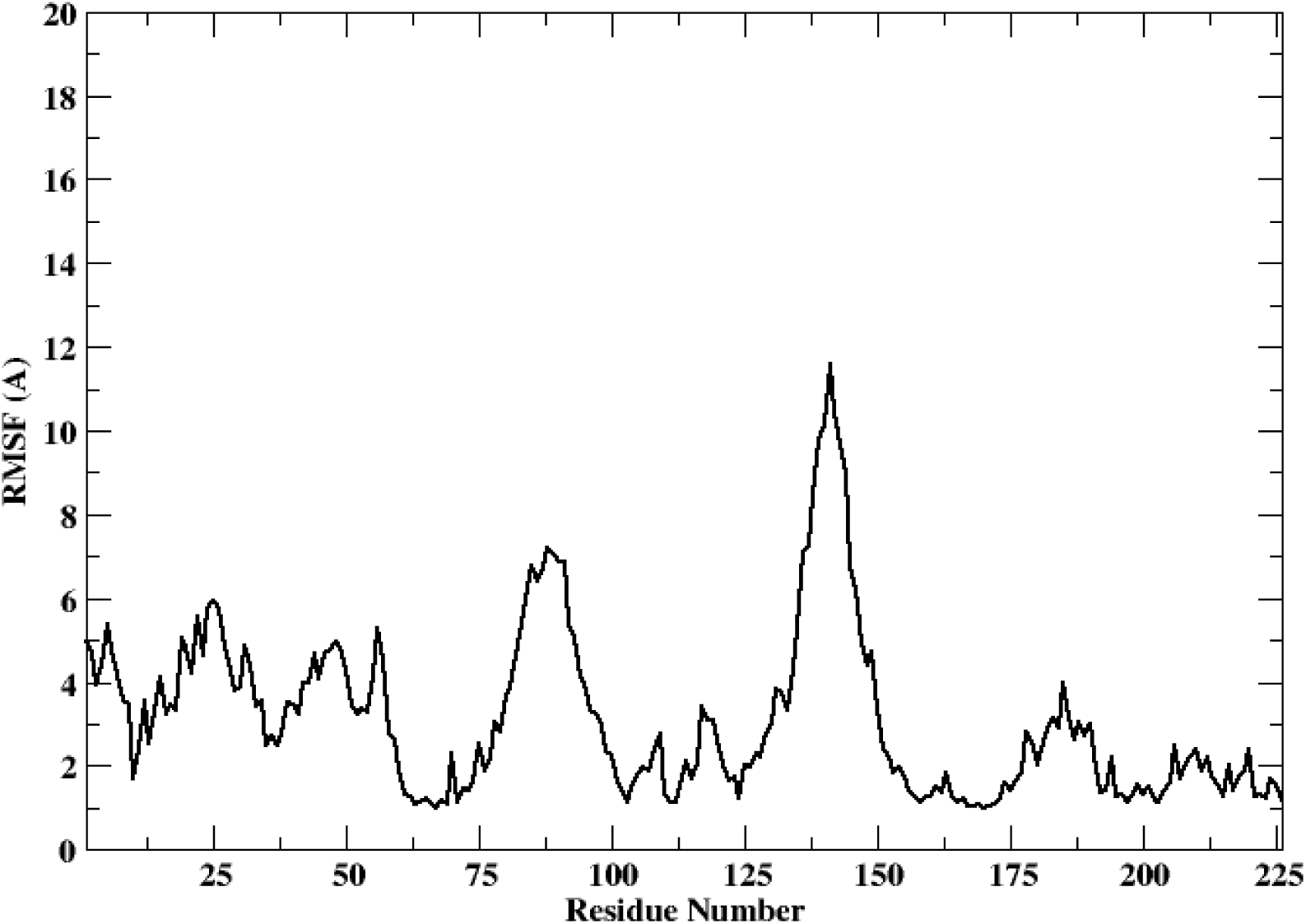
RMSF of the docked vaccine construct

GetContact tool was used for the calculation of hydrogen bonds and Van der Waal’s interactions between vaccine construct and TLR2 - TLR6. All the interactions having occupancy of ≥ 20% have been shown in **Supplementary Tables (2,3) and Supplementary Figures (2,3).** TLR2 forms eleven hydrogen bonds with the vaccine construct. Among these, four hydrogen bonds are between the epitope regions and TLR2, and the remaining seven hydrogen bonds are between the linker region or β-defensin region and TLR2. Besides hydrogen bonds, the vaccine construct formed thirty-eight Van der Waal’s interactions with TLR2. Of these, twenty-six interactions were between epitope regions of the vaccine construct and TLR2. Similarly, TLR6 formed twenty-six hydrogen and sixty-two Van der Waal’s interactions with the vaccine construct. Of the twenty-six hydrogen bonds, seven were between epitope regions and TLR6; fourteen were between the adjuvant and the TLR6. Similarly, of the sixty-two Van der Waal’s interactions, twenty-five involved epitope regions. Thus, the TLR6 receptor tends to form the maximum number of interactions with vaccine construct as compared to TLR2.

Further, the free energy of binding between different components of the complex, namely, vaccine construct-TLR2/TLR6, vaccine construct and TLR2-TLR6 complex, and between TLR2-TLR6, were computed and are shown in **Figure 8**. It can be observed that the vaccine construct shows strong free energy of binding with TLR2-TLR6 complex with values around -80 to -100 kcal/mol. Among the two TLRs, the vaccine construct showed better binding free energy with TLR6 as compared to TLR2. The vaccine construct and TLR2 binding free energy lies in the range of -25 to -27 kcal/mol, while it lies in the range of -60 to -75 kcal/mol for the vaccine construct and TLR6. The free energy of binding between two TLRs lies in the range of 0 to 5 kcal/mol, which indicates weak binding.

**Figure 8:**
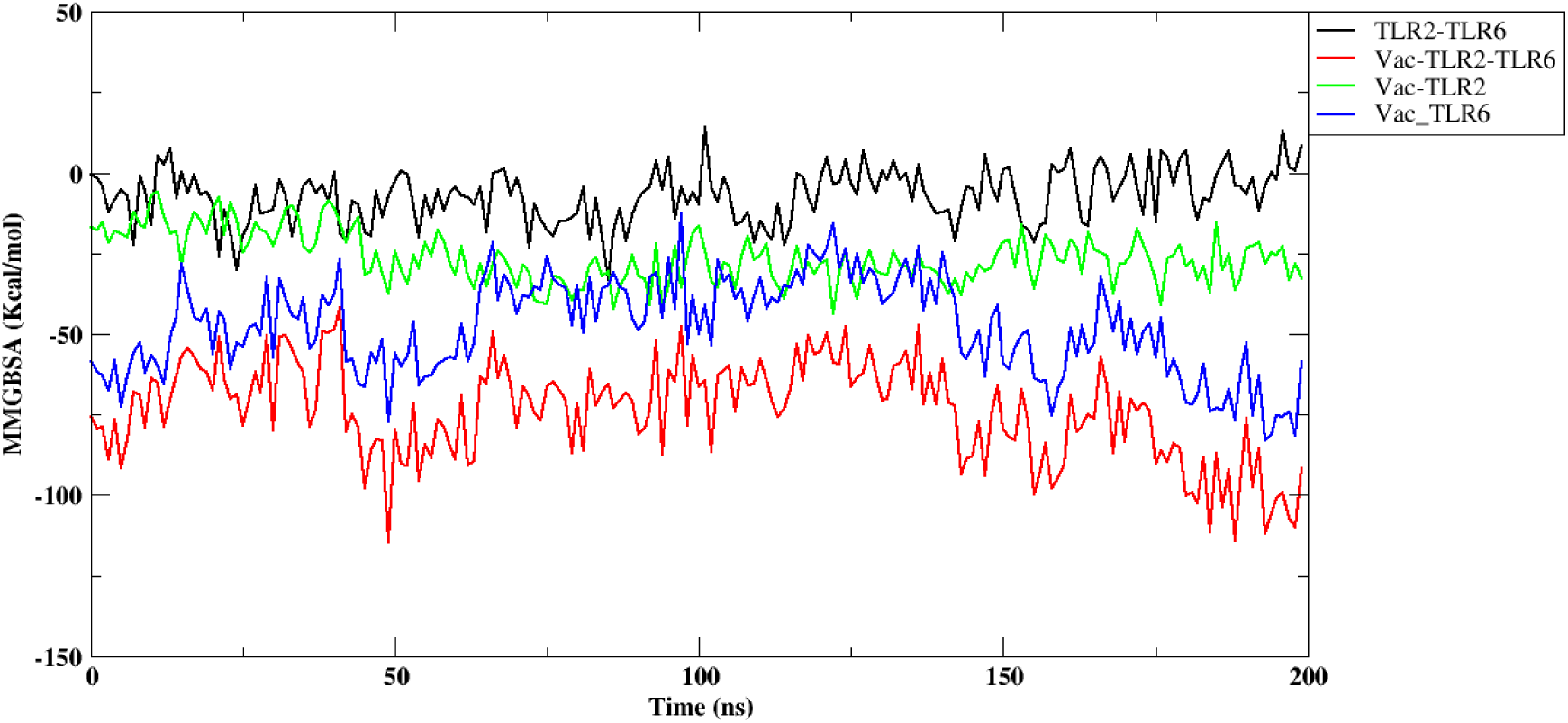
Free energy of binding between vaccine construct and TLR2-TLR6 complex. TLR2 and TLR6 (black), vaccine construct and TLR2-TLR6 complex (red), vaccine construct and TLR2 (green), vaccine construct and TLR6 (blue).

### 3.9 Codon optimization and In silico cloning

Vector Builder was used for codon optimization in the *Escherichia coli* (strain K12) expression system. The GC content was 59.43% and Codon Adaptation Index (CAI) was 0.95. A codon-optimized sequence of vaccine construct was used for *in-silico* insertion into pET30b (+)between XhoI and NdeI restriction sites using the SnapGene tool **(Figure 9).**

**Figure 9:**
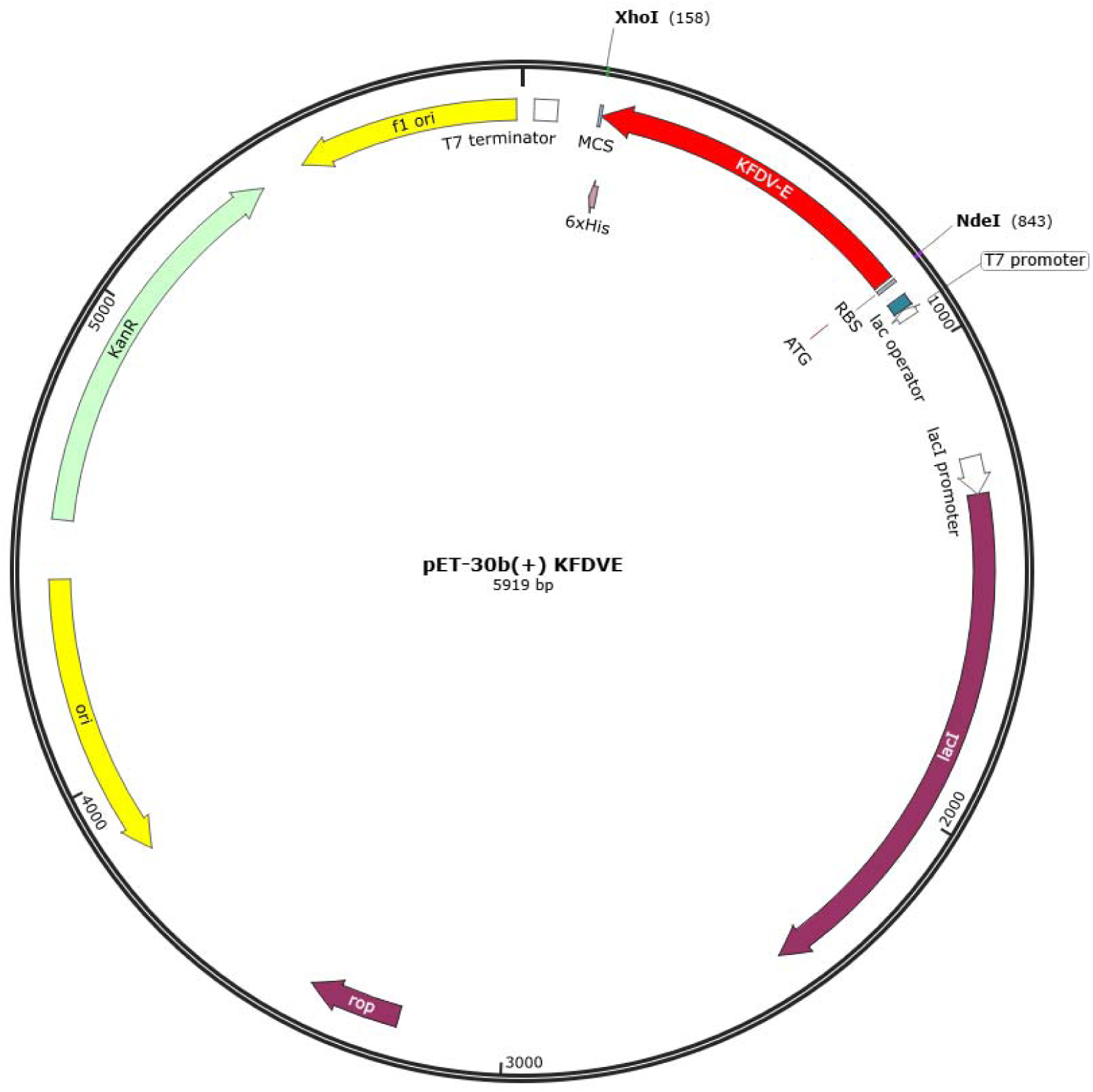
*In-silico* cloning of multi-epitope vaccine in pET30b(+) using Snapgene tool. The red color part represents the epitope vaccine sequence inserted between NdeI and XhoI.

## Discussion

Kyasanur Forest Disease is spreading to newer areas. Vaccination has been one of the primary preventive measures. However, the efficacy of the existing vaccine may vary, and it is crucial to boost its effectiveness or develop new vaccines with improved efficacy. A highly immunogenic vaccine is required to fight against this disease and its rapid dispersal to newer areas. The availability of genomic data and immunoinformatics tools helps develop new epitope vaccines and screen these vaccine candidates in less time. Many effective peptide vaccines against infectious diseases developed by employing the reverse vaccinology approaches and the immunoinformatic tools were experimentally validated^48^.

Different research groups followed computational vaccine design methods for predicting B-cell and T-cell epitopes to enhance the speed of vaccine development^20–22^. We have analyzed genome sequences from 1957-2022, and it was noted that there is no evidence of genetic recombination among the isolates.

In this study, the envelope (E) protein of KFDV was chosen to prepare a multi-epitope vaccine candidate. Previous studies conducted on KFDV showed the suitability of the envelope protein as a candidate for newer vaccines and diagnostics^21^. The envelope protein is responsible for the entry of the virus into the host cells by receptor binding and fusion of viral and cellular membranes^8^. The Ecto-domain-III (EIII) of the envelope protein of flaviviruses forms specific neutralizing epitopes^49^. In earlier studies on immune response, a notable B- and T-cell response was observed in KFDV-infected patients^50^. This suggests the need to develop a potential vaccine for KFD that can elicit both humoral and cellular immune responses. Different immunoinformatics tools are used for the prediction and prioritization of a set of B-cell, MHC I, and MHC II epitopes. MHC class I and MHC class II epitopes of envelope protein were predicted using HLA reference alleles. MHC molecules present the peptide epitopes to the T-cell receptors (TCR). MHC I present the peptides to CTL through the cytosolic pathway and MHC II to the HTL through the endocytic pathway. The non-allergen, non-toxic, and IFN-positive epitopes were selected for the final vaccine construct. We designed the multi-epitope vaccine construct by joining the selected predicted epitopes using linkers. There are many *in-silico* multi-epitope vaccine candidates for KFDV and other infectious diseases^22,51–53^. The glycosylation pattern of the epitope has effects on the immunogenicity of the vaccine^54,55^. The prioritized epitopes were compared with the key amino acids of the envelope protein. One of the selected epitopes 57-REYCLHAKLANSKVAARCPAM-77 contains the Asp67. The N-glycosylation of the Asp67 of E-protein is responsible for the Dengue viral assembly or exit and dendritic cell infection by interacting with DC-SIGN receptors^56^. The same Asp67 residue was present in the E-protein of KFDV in the studies conducted by Dey et al., suggesting a similar mechanism of viral recognition and infection in KFD^21^. The ecto-domain-III (EIII) contains the receptor binding region of the envelope protein. The amino acids that form the receptor binding regions are K315, L388, H390, Q391, K395, and F398^21^. The epitope 384-YVGELSHQW-392 present in the vaccine construct also contains the L388, H390, and Q391 that will enhance the receptor binding ability of the multi-epitope vaccine. Usually, the purified proteins or peptides are less immunogenic, so there is a need to add an adjuvant at the terminal of the peptide to enhance immunogenicity. To increase the immunogenicity of the purified protein β-defensin adjuvant was added ^41^ at the N-terminal end along with His tag at the C-terminal end for purification for the protein. Toll-like receptors (TLR) recognize the microbial components and elicit immune responses by inducing innate immunity followed by adaptive immunity^57–59^. The multi-epitope vaccine construct was docked with the TLR2, TLR6, and TLR2-TLR6 complex. The RMSD of the docked complex (vaccine with TLR2-TLR6 complex) was around 0.6 +/- 0.2, indicating a good quality model. Many studies have reported the role of TLR2 as a host receptor for the envelope protein to evoke the innate immune response^59^. TLR2 is the most ubiquitous and is the only TLR that can form heterodimers with more than two other types of TLRs. It forms a heterodimer with TLR6. It interacts with a large number of other non-TLR molecules, thereby increasing its capacity to recognize pathogen-associated molecular patterns (PAMPs)^60^. TLR2 also interacts with molecules like human β-defensin-3, heat shock proteins, and high mobility group box 1 protein and considers them as endogenous ligands^61^. We have used the β-defensin as an adjuvant, which can be recognized by TLR2-TLR1 and TLR2-TLR6 heterodimers, increasing the interactive ability of the vaccine construct with the TLR. The heterodimerization increases the range of recognizable motifs by the receptors. Four hydrogen bonds are formed between the epitope regions and TLR2, and the remaining seven hydrogen bonds are formed between the linker region or β-defensin region and TLR2. In the case of vaccine construct with the TLR6, more interactions with the adjuvant region were observed, resulting in more stability. Overall, the vaccine construct forms a stable complex with TLR2-TLR6 compared to TLR2 alone. In most of the previous studies of the KFDV multi-epitope vaccine, molecular docking was done with TLR2 alone^20,22^, which is a major limitation as TLR2 recognizes the viral proteins in their heterodimeric forms^62^. In this study, the ability of the TLR2 to form a heterodimer with the TLR6 was also considered, and the vaccine construct was docked with the TLR2-TLR6 complex. The top-scoring docked model of the vaccine obtained was with the TLR2-TLR6 complex. The free energy of binding between two TLRs lies in the 0 to 5 kcal/mol range, indicating weak binding. The vaccine construct has a strong free energy of binding with the TLR2-TLR6 complex with values around -80 to -100 kcal/mol, showing the maximum stable interaction between the vaccine construct and the TLR complex.

Codon optimization was done to enhance the translation efficiency in *E. coli* (Strain K12). The GC content of 59.43% (30-70%) and CAI 0.95 (0.8-1.0) obtained were in the optimum range for protein expression in the target organism.

Constructing an immunoinformatics-based multi-epitope vaccine against KFDV represents a proactive approach to addressing this emerging infectious disease. Further research, including *in-vitro* and *in-vivo* studies, will be necessary to validate the vaccine’s efficacy, safety, and potential for clinical use. In conclusion, an immunoinformatics-based multi-epitope vaccine targeting KFDV holds significant promise in the fight against KFD.

## Limitations of the study

It needs to be mentioned that in the present study, only the major antigenic protein, namely envelope protein, was used in the design of multi-epitope vaccine candidates. The inclusion of other structural (capsid and membrane) and non-structural (NS1-NS5) viral proteins would provide a more comprehensive immune response. The placement of predicted B- and T-cell epitopes in the vaccine construct is also known to play a role in eliciting the desired immunogenicity; hence, different arrangements of the epitopes need to be verified. Prediction of MHC-I/II and their binding with epitopes promises to provide a more rigorous criterion for epitope prioritization and hence is proposed to be carried out in future studies. Molecular docking studies of the vaccine construct with other TLRs would also enable a comparative analysis of the binding energies.

## Financial support & sponsorship

The grant was provided by the Indian Council of Medical Research, New Delhi, India under the extramural project ‘Sustainable laboratory network for monitoring of Viral Hemorrhagic Fever viruses in India and enhancing bio-risk mitigation for High risk group pathogen’ with the grant number: VIR/28/2020/ECD-1 dated 10.05.2023. The funders had no role in study design, data collection and analysis, decision to publish, or preparation of the manuscript.

## Acknowledgment

Authors gratefully acknowledge the encouragement and support extended by Dr. Sheela Godbole, Director In-Charge, ICMR-National Institute of Virology, Pune. We also acknowledge the excellent support from Dr. Abhinendra Kumar, Mr. Rajen Lakra, Mr. Prasad Sarkale, Mr. Hitesh Dighe, Mr. Deepak Mali, Ms. Pranita Gawande, Ms. Ujjwala Gaikwad, Ms. Kumari Vaishnavi and Mrs. Pratiksha Vedapathak of the Maximum Containment Facility of ICMR-National Institute of Virology Pune.

## Notes

### Competing Interest Statement

The authors have declared no competing interest.

