## Supplement for "Construction of an Immunoinformatics-Based Multi-Epitope Vaccine Candidate targeting Kyasanur Forest Disease Virus"

**Supplementary Figures and Tables**

**List of supplementary Figures**

**S Figure 1:** Ramachandran plot

**S Figure 2:** Interaction plot for all the hydrogen bonds and Van der Waal’s interactions in the docked complex of vaccine construct with TLR2 at different time frames of simulations.

**S Figure 3:** Interaction plot for all the hydrogen bonds and Van der Waal’s interactions in the docked complex of vaccine construct with TLR6 at different time frames of simulations.

**List of supplementary Tables**

**S Table 1:** Representative isolates identified based on the lineages observed in the phylogenetic tree**.**

**S Table 2:** Table showing hydrogen and Van der Waal’s interacting residues between TLR2 receptor and vaccine construct

**S Table 3:** Table showing hydrogen and Van der Waal’s interacting residues between TLR6 receptor and vaccine construct

**Supplementary Figures**

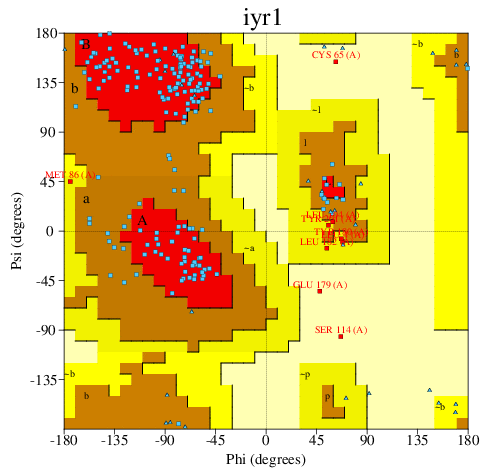

**S Figure 1:** Ramachandran plot

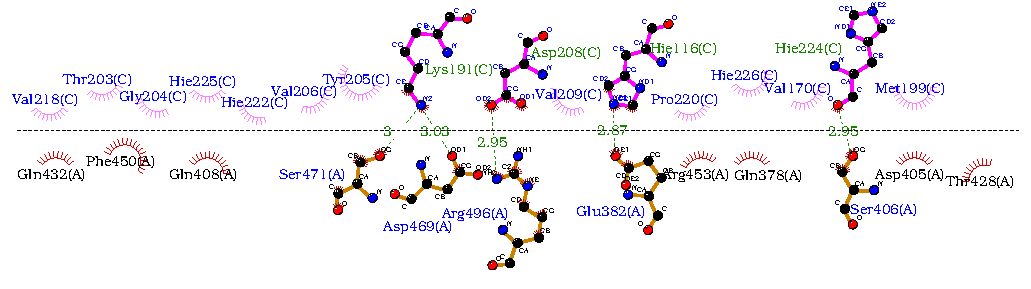

*50ns*

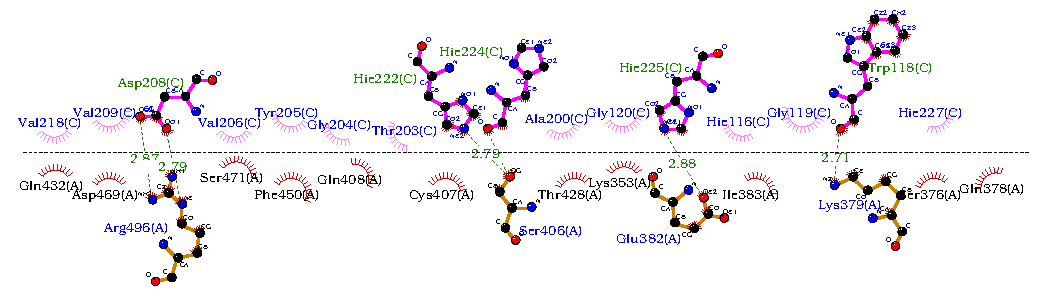

*100ns*

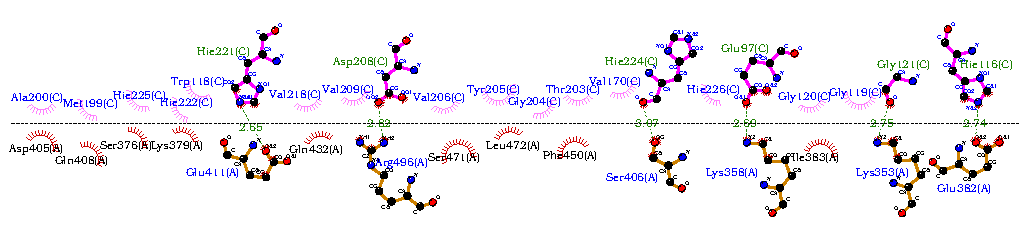

*150ns*

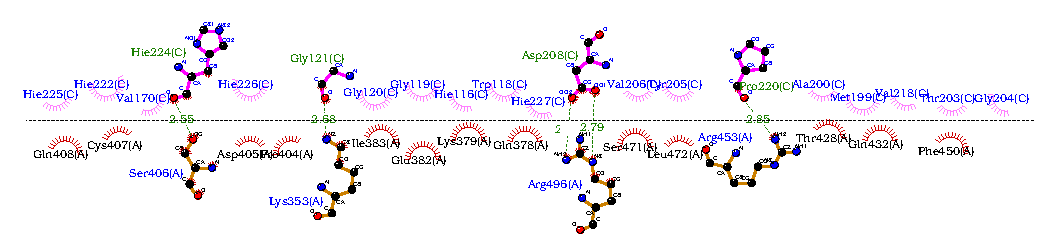
*200ns*

**S Figure 2**: Interaction plot for all the hydrogen bonds and Van der Waal’s interactions in the docked complex of vaccine construct with TLR2 at different time frames of simulations.

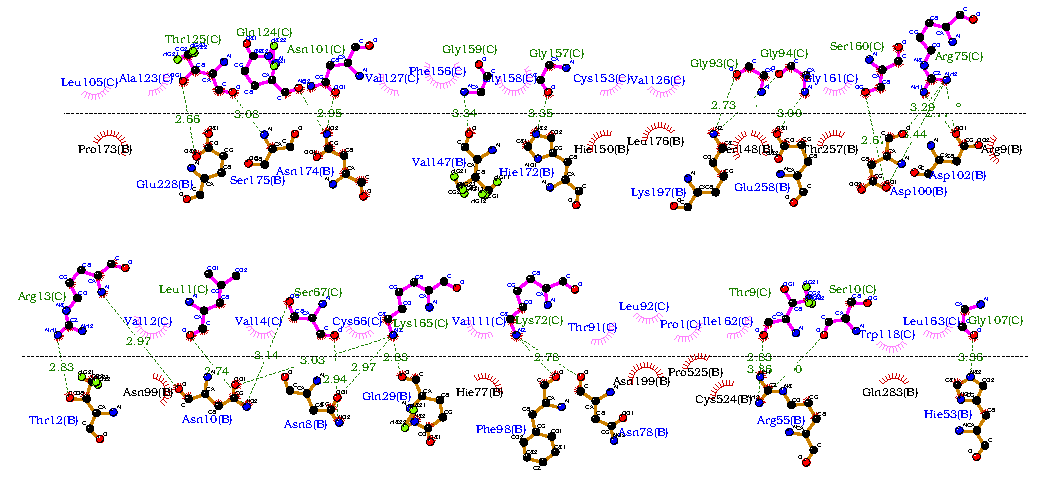

*50ns*

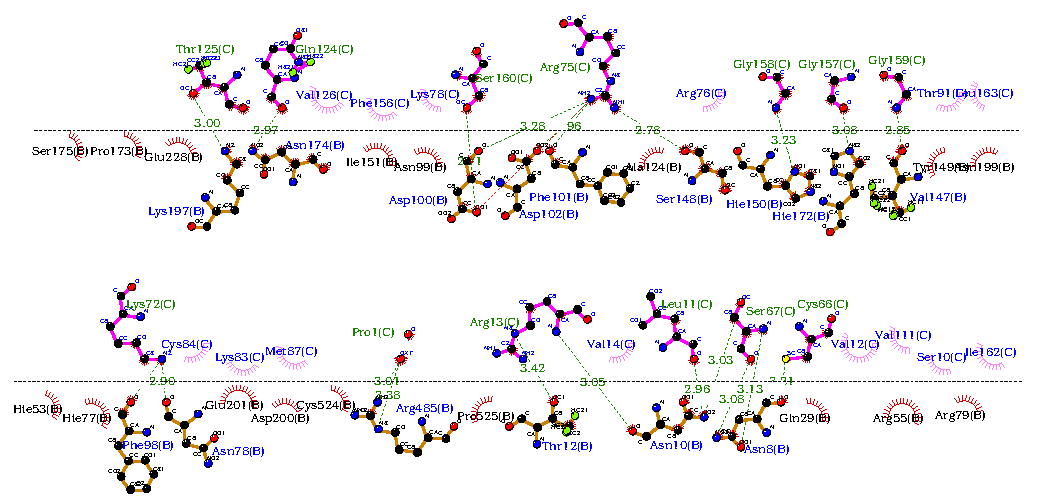

*100ns*

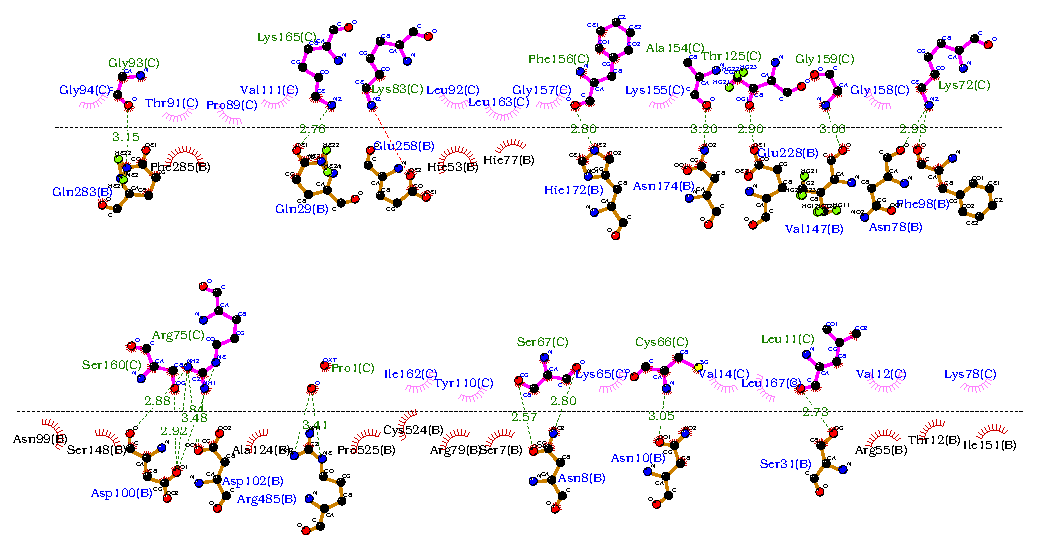

*150ns*

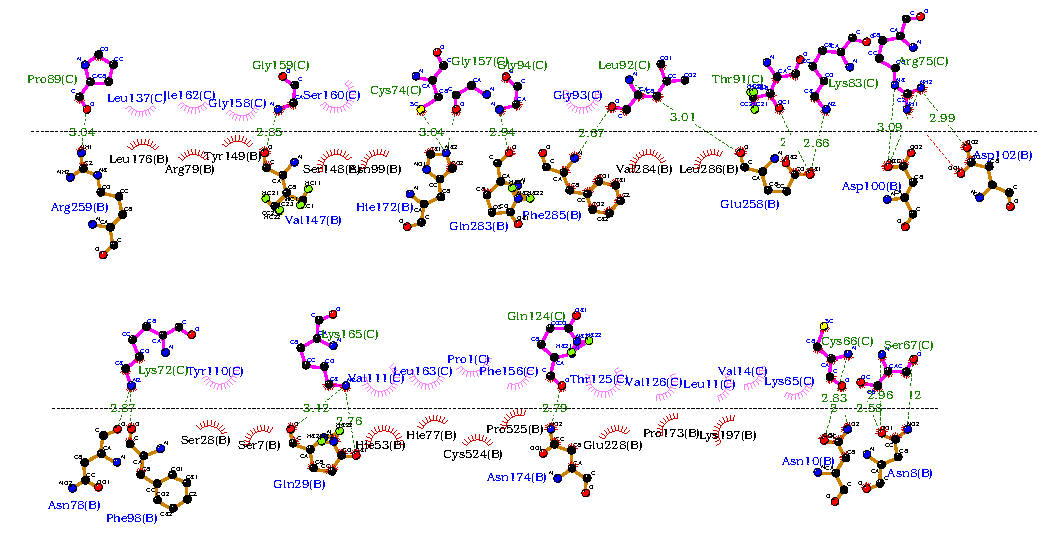
*200ns*

**S Figure 3**: Interaction plot for all the hydrogen bonds and Van der Waal’s interactions in the docked complex of vaccine construct with TLR6 at different time frames of simulations.

**Supplementary Tables**

| **Representative sequence** | **Lineage details** |
| --- | --- |
| P9605_P1-KA-H-1957 | L 1 |
| MG720114_NIV_1722297-KA-H-2017 | L2.2.1 |
| OR162010_MCL-19-H-561-MH-H-2019 | L2.2.2 |
| MG720091_NIV_164187(1297)-KA-H-2016 | L 2.1 |
| OR162020_MCL-19-H-1180-MH-H-2019 | L2.2.2 |
| MG720116_MCL-17-T-296-GA-2017 | L2.2.2 |
| OR179916_MCL-20-H-690-MH-H-2020 | L2.2.2 |
| OR162012_MCL-19-H-553-MH-H-2019 | L2.2.2 |
| OR162018_MCL-19-H-897-MH-H-2019 | L2.2.2 |
| MG720114-NIV_1722297-KA-H-2017 | L2.2.1 |

**S Table 1**: Representative isolates identified based on the lineages observed in the phylogenetic tree

| **Hydrogen Bond** | | | | **Van der Waals** | | | |
| --- | --- | --- | --- | --- | --- | --- | --- |
| **TLR2_Residue** | **Vaccine Construct Residue** | **Occupancy** |  | | **TLR2_Residue** | **Vaccine Construct Residue** | **Occupancy** |
| GLN378 | HIE225 | 0.207 |  |  | PRO404 | HIE225 | 0.21 |
| GLN378 | HIE226 | 0.816 |  |  | GLU411 | HIE220 | 0.22 |
| *LYS379* | *TRP117* | *0.222* |  |  | *LYS358* | *GLU96* | *0.22* |
| GLU382 | GLY119 | 0.469 |  |  | *GLN432* | *GLU218* | *0.221* |
| *GLU382* | *HIE115* | *0.935* |  |  | *PHE450* | *THR202* | *0.266* |
| *ARG496* | *ASP207* | *1* |  |  | *ASP405* | *MET198* | *0.274* |
| *LYS358* | *GLU96* | *0.228* |  |  | *LEU472* | *TYR204* | *0.284* |
| LYS353 | GLY118 | 0.386 |  |  | ILE383 | GLY119 | 0.35 |
| LYS353 | GLY120 | 0.584 |  |  | *GLN432* | *PRO219* | *0.36* |
| SER406 | HIE221 | 0.528 |  |  | *SER376* | *TRP117* | *0.37* |
| SER406 | HIE223 | 0.766 |  |  | GLN378 | HIE225 | 0.428 |
|  | | |  |  | *SER471* | *ASP207* | *0.449* |
|  |  |  |  |  | GLU382 | GLY118 | 0.451 |
|  |  |  |  |  | *THR428* | *MET198* | *0.457* |
|  |  |  |  |  | LYS353 | GLY119 | 0.476 |
|  |  |  |  |  | ILE383 | GLY118 | 0.501 |
|  |  |  |  |  | *GLN432* | *VAL217* | *0.503* |
|  |  |  |  |  | GLU382 | GLY119 | 0.539 |
|  |  |  |  |  | *ARG496* | *VAL208* | *0.539* |
|  |  |  |  |  | *GLN378* | *TRP117* | *0.559* |
|  |  |  |  |  | SER406 | HIE225 | 0.563 |
|  |  |  |  |  | LYS353 | GLY118 | 0.592 |
|  |  |  |  |  | CYS407 | HIE221 | 0.602 |
|  |  |  |  |  | THR428 | ALA199 | 0.62 |
|  |  |  |  |  | *SER406* | *VAL169* | *0.626* |
|  |  |  |  |  | LYS353 | GLY120 | 0.628 |
|  |  |  |  |  | SER406 | HIE221 | 0.641 |
|  |  |  |  |  | *SER471* | *VAL205* | *0.685* |
|  |  |  |  |  | GLN408 | HIE224 | 0.702 |
|  |  |  |  |  | *PHE450* | *TYR204* | *0.751* |
|  |  |  |  |  | GLN408 | HIE221 | 0.763 |
|  |  |  |  |  | *LYS379* | *TRP117* | *0.786* |
|  |  |  |  |  | GLN378 | HIE226 | 0.822 |
|  |  |  |  |  | *GLU382* | *HIE115* | *0.928* |
|  |  |  |  |  | SER406 | HIE223 | 0.938 |
|  |  |  |  |  | *SER471* | *TYR204* | *0.948* |
|  |  |  |  |  | *PHE450* | *GLY203* | *0.971* |
|  |  |  |  |  | *ARG496* | *ASP207* | *1* |

**S Table 2**: Table showing hydrogen and Van der Waal’s interacting residues between TLR2 receptor and vaccine construct

| **Hydrogen Bond** | | |  | **Van der Waals** | | |
| --- | --- | --- | --- | --- | --- | --- |
| **TLR6_Residue** | **Vaccine Construct Residue** | **Occupancy** |  | **TLR6_Residue** | **Vaccine Construct Residue** | **Occupancy** |
| *ASN174* | *THR124* | *0.2* |  | ARG55 | LEU10 | 0.201 |
| PHE101 | ARG74 | 0.207 |  | SER31 | VAL11 | 0.207 |
| *ARG259* | *ALA89* | *0.225* |  | HIE150 | GLY157 | 0.211 |
| GLN283 | GLY93 | 0.262 |  | *VAL284* | *LEU91* | *0.227* |
| *PHE285* | *LEU91* | *0.269* |  | *SER7* | *TYR109* | *0.243* |
| *GLU258* | *LEU91* | *0.278* |  | PHE101 | ARG74 | 0.252 |
| ASN10 | SER66 | 0.288 |  | *ARG259* | *ALA89* | *0.252* |
| THR12 | ARG12 | 0.322 |  | *GLU258* | *THR90* | *0.26* |
| GLU258 | LYS82 | 0.363 |  | GLU258 | GLY92 | 0.262 |
| ASN10 | LEU10 | 0.367 |  | *SER175* | *THR124* | *0.269* |
| ILE151 | LYS77 | 0.372 |  | *HIE53* | *LYS164* | *0.279* |
| SER31 | LEU10 | 0.431 |  | HIE150 | ARG74 | 0.296 |
| SER148 | ARG74 | 0.455 |  | HIE172 | CYS73 | 0.298 |
| ASN10 | ARG12 | 0.491 |  | *ARG55* | *ILE161* | *0.3* |
| ASP102 | ARG74 | 0.578 |  | *PHE285* | *LEU91* | *0.302* |
| ASN10 | CYS65 | 0.635 |  | *GLU258* | *LEU91* | *0.306* |
| HIE172 | GLY156 | 0.67 |  | HIE77 | LYS71 | 0.329 |
| PHE98 | LYS71 | 0.701 |  | GLN283 | GLY92 | 0.334 |
| ASP100 | SER159 | 0.783 |  | GLU258 | LYS82 | 0.352 |
| *ASN174* | *GLN123* | *0.807* |  | GLN29 | SER66 | 0.355 |
| *GLN29* | *LYS164* | *0.846* |  | ARG55 | VAL11 | 0.359 |
| ASP100 | ARG74 | 0.874 |  | SER148 | GLY158 | 0.363 |
| ASN78 | LYS71 | 0.9 |  | *ARG79* | *ILE161* | *0.368* |
| VAL147 | GLY158 | 0.949 |  | *HIE172* | *PHE155* | *0.386* |
| *GLU228* | *THR124* | *0.975* |  | GLN283 | GLY93 | 0.389 |
| ASN8 | SER66 | 0.996 |  | ALA124 | ARG74 | 0.392 |
|  | | |  | *GLN29* | *VAL110* | *0.407* |
|  |  |  |  | TYR149 | GLY157 | 0.408 |
|  |  |  |  | *HIE77* | *LEU162* | *0.428* |
|  |  |  |  | *ASN8* | *VAL110* | *0.445* |
|  |  |  |  | THR12 | VAL11 | 0.447 |
|  |  |  |  | SER31 | LEU10 | 0.484 |
|  |  |  |  | SER148 | ARG74 | 0.485 |
|  |  |  |  | THR12 | ARG12 | 0.486 |
|  |  |  |  | ASN10 | ARG12 | 0.512 |
|  |  |  |  | ASP102 | ARG74 | 0.545 |
|  |  |  |  | ASN10 | VAL13 | 0.576 |
|  |  |  |  | *ASN174* | *PHE155* | *0.609* |
|  |  |  |  | SER148 | GLY157 | 0.613 |
|  |  |  |  | ASN10 | SER66 | 0.617 |
|  |  |  |  | *LYS197* | *VAL125* | *0.631* |
|  |  |  |  | ILE151 | LYS77 | 0.642 |
|  | | |  | *LYS197* | *THR124* | *0.673* |
|  |  |  |  | *PRO173* | *THR124* | *0.733* |
|  |  |  |  | ASN10 | CYS65 | 0.741 |
|  |  |  |  | SER148 | SER159 | 0.768 |
|  |  |  |  | *HIE172* | *GLY156* | *0.784* |
|  |  |  |  | ASN99 | SER159 | 0.816 |
|  |  |  |  | *ASN174* | *GLN123* | *0.842* |
|  |  |  |  | ASN10 | LEU10 | 0.861 |
|  |  |  |  | *HIE53* | *LEU162* | *0.868* |
|  |  |  |  | ASP100 | ARG74 | 0.873 |
|  |  |  |  | VAL147 | GLY157 | 0.895 |
|  |  |  |  | *ASN174* | *THR124* | *0.903* |
|  |  |  |  | ASN8 | CYS65 | 0.906 |
|  |  |  |  | ASP100 | SER159 | 0.959 |
|  |  |  |  | *GLN29* | *LYS164* | *0.972* |
|  |  |  |  | VAL147 | GLY158 | 0.979 |
|  |  |  |  | *GLU228* | *THR124* | *0.995* |
|  |  |  |  | ASN8 | SER66 | 0.998 |
|  |  |  |  | PHE98 | LYS71 | 0.998 |
|  |  |  |  | ASN78 | LYS71 | 1 |

**S Table 3**: Table showing hydrogen and Van der Waal’s interacting residues between TLR6 receptor and vaccine construct
